# Putative host-derived insertions in the genomes of circulating SARS-CoV-2 variants

**DOI:** 10.1101/2022.01.04.474799

**Authors:** Yiyan Yang, Keith Dufault-Thompson, Rafaela Salgado Fontenele, Xiaofang Jiang

**Author notes:** Correspondence: Xiaofang Jiang.

## Abstract

Insertions in the SARS-CoV-2 genome have the potential to drive viral evolution, but the source of the insertions is often unknown. Recent proposals have suggested that human RNAs could be a source of some insertions, but the small size of many insertions makes this difficult to confirm. Through an analysis of available direct RNA sequencing data from SARS-CoV-2 infected cells, we show that viral-host chimeric RNAs are formed through what are likely stochastic RNA-dependent RNA polymerase template switching events. Through an analysis of the publicly available GISAID SARS-CoV-2 genome collection, we identified two genomic insertions in circulating SARS-CoV-2 variants that are identical to regions of the human 18S and 28S rRNAs. These results provide direct evidence of the formation of viral-host chimeric sequences and the integration of host genetic material into the SARS-CoV-2 genome, highlighting the potential importance of host-derived insertions in viral evolution.

**IMPORTANCE:** Throughout the COVID-19 pandemic, the sequencing of SARS-CoV-2 genomes has revealed the presence of insertions in multiple globally circulating lineages of SARS-CoV-2, including the Omicron variant. The human genome has been suggested to be the source of some of the larger insertions, but evidence for this kind of event occurring is still lacking. Here, we leverage direct RNA sequencing data and SARS-CoV-2 genomes to show host-viral chimeric RNAs are generated in infected cells and two large genomic insertions have likely been formed through the incorporation of host rRNA fragments into the SARS-CoV-2 genome. These host-derived insertions may increase the genetic diversity of SARS-CoV-2 and expand its strategies to acquire genetic materials, potentially enhancing its adaptability, virulence, and spread.

## INTRODUCTION

During the COVID-19 pandemic, insertions have been frequently acquired in SARS-CoV-2 lineages (1–4). Insertions have been associated with several globally circulating lineages, including the insertion of one amino acid at position 146 of the S protein (ins146N) of the variant of interest Mu (B.1.621) (4), insertions at the recurrent insertion site 214 of the NTD region on the S protein that occurred in the lineages B.1.214.2 (ins214TDR) and A.2.5 (ins214AAG) (1), and the insertion ins214EPE in the recently-emerged variant of concern Omicron (5). Although there is insufficient evidence to show the direct impact these insertions have on viral spread and interference with immune responses, the fact that variants carrying those insertions have circulated for long periods suggests that they might be advantageous or neutral for the transmission. Results from a long-term *in vitro* experiment where SARS-CoV-2 was co-incubated with highly neutralizing antibodies have also shown that an 11 amino acid insertion (ins248KTRNKSTSRRE) at the NTD N5 loop of the S protein was able to drive antibody escape suggesting a potential role of insertions in enhancing infectivity and virulence (6). Taken together, insertions have the potential to increase genetic diversity in SARS-CoV-2 and contribute to the continued evolution of the virus.

Previous research has shown that most small insertions in the SARS-CoV-2 genome likely originated from template sliding, local duplication, or template switching between viruses (2). Longer insertions (equal or larger than nine nucleotides) have been detected in multiple coronavirus genomes, including in variants of concern like the Omicron variant, but their origin remains unknown. Host genetic material has been suggested as a possible source for these insertions (5, 7). Venkatakrishnan et al. suggested that the unique insertion (ins214EPE) in the Omicron variant could have originated from the human common cold virus HCoV-229E or the human genome based on BLAST search (5), and the human genome has been speculated to be the source of multiple other small insertions (7). However, given that these insertion sequences are typically short, sequence comparisons tend to be less informative, and false-positive matches have a high chance of occurring. Additionally, coronavirus replication occurs in modified endoplasmic reticulum-derived double-membrane vesicles, providing a physical barrier between viral and host genetic material (8), and coronavirus replication complexes are known to contain enzymes with proofreading activity (9), both of which likely play roles in limiting the formation of host-virus chimeric sequences.

Human-derived insertions in the SARS-CoV-2 genome would likely be generated through RdRp-driven template switching events between SARS-CoV-2 and host mRNA. While template switching events between coronaviruses are common (10–13) and likely contribute to the emergence of SARS-CoV-2 lineages including the deltacron variant (14), template switching events between coronaviruses and host RNAs are rarely documented (15, 16). Chimeric reads between SARS-CoV-2 RNA and human RNA have been detected but were interpreted as a signal of SARS-CoV-2 integration into the human genome in a previous controversial study (17). Others have suggested that the chimeric reads were likely to be template switching artifacts mediated by reverse transcriptase or PCR during library preparation (18–21). One possible explanation that was largely omitted in these studies is that the SARS-CoV-2-host chimeric RNA could be generated by RdRp-driven template switching.

Here, to investigate the possible existence of SARS-CoV-2-host chimeric RNA, we take advantage of the publically available Nanopore direct RNA sequencing data of SARS-CoV-2. Direct RNA-seq sequences the individual polyadenylated RNAs directly mitigating the possible formation of chimeric reads during library preparation or amplification. We first identified SARS-CoV-2-host chimeric RNA from direct RNA-seq data and showed that RdRp-driven template switching between SARS-CoV-2 and host mRNA occurs, but it is infrequent and stochastic. We also found that highly expressed host genes and structural RNA genes have a higher chance to be observed in chimeric RNA reads. We then systematically analyzed the SARS-CoV-2 genomes deposited in the GISAID database (22), resulting in the identification of two insertions in functional SARS-CoV-2 genomes that likely originated from the host 18S and 28S rRNAs.

## RESULTS

### Host-virus mRNA chimera are rare but do exist

We first analyzed direct RNA-seq data from SARS-CoV-2 infected cell lines to identify sequences formed from chimeric host-viral RNAs. The direct RNA-seq data were quality filtered and mapped to both the host and SARS-CoV-2 transcriptomes to identify potential chimeric sequences. Out of the 30 samples that were analyzed, host-viral chimeric reads were detected in 16 of the samples with an average of 0.029% (standard deviation 0.048%) of the reads mapped to SARS-CoV-2 being chimeric (Supplementary Table 1). Chimeric reads were typically rare, making up 0.206% of one sample, but less than 0.06% of the other 15 samples, and these rates may be an overestimation due to the cell lines used compared to what would be observed in *in vivo* conditions. Additionally, chimeric reads detected in five samples were further investigated using paired-end sequencing short reads from the same samples (Supplementary Table 2). Approximately 1.4% (5 out of 357) of chimeric reads were supported by at least five read pairs spanning the junctions. This finding implies that a small fraction of the host-viral chimeric mRNA molecules could function as templates for RNA replication.

We then analyzed the chimeric reads to identify trends in how the viral and host RNA sequences were joined. All the viral-derived sequences in chimeric reads were annotated as positive-sense RNA and a majority (92.49%) of the reads contained host-derived positive-sense sequences. Upon further examination, the few host reads that were identified as being negative-sense were largely long non-coding RNAs that were present in the raw reads as the negative-sense sequences, making it likely that they were mis-annotated rather than actually being derived from negative-sense RNA. These results suggest that the host-viral chimeric sequences are not the result of the integration of the viral genetic material into the host genome, which would have resulted in a nearly equal mix of positive and negative sense viral sequences (17). Most likely, these host-viral chimeric sequences were created from positive-to-positive-strand template switching events (23, 24).

### Viral-host chimeric read formation is likely a stochastic process

The chimeric reads were then analyzed to determine if there were any patterns in the composition of the sequences and in which positions relative to the references they were formed. Both viral to host and host to viral chimeric sequences were detected in the direct RNA-seq data, but the chimeric reads did not show a preference for either organization (Supplementary Table 1). Both types of sequences were seen in approximately the same frequency, with viral to host reads making up 55% of the chimeric sequences and host to viral reads making up 45%. This lack of strong preference may indicate that host RNA can be readily recognized by viral RdRp, but other factors like the exclusion of host RNA by the formation of the double-membrane vesicles might prevent the formation of chimeric RNAs. When examining the positions of the junctions on the viral RNA sequences, we found there was a bias toward the junction sites being located in the dense coding region near the three prime end of the sequence, with fewer junctions being identified in the ORF1ab genes, the largest region of the genome (Fig.1). This is likely due to the ORF1ab region not being retained in the canonical SARS-CoV-2 subgenomic RNAs resulting in fewer viral RNAs being synthesized with these regions that could form chimeric RNAs (25). It suggests that the process by which chimeric sequences are formed is likely stochastic, depending on the availability of template RNA molecules.

**Fig.1.**
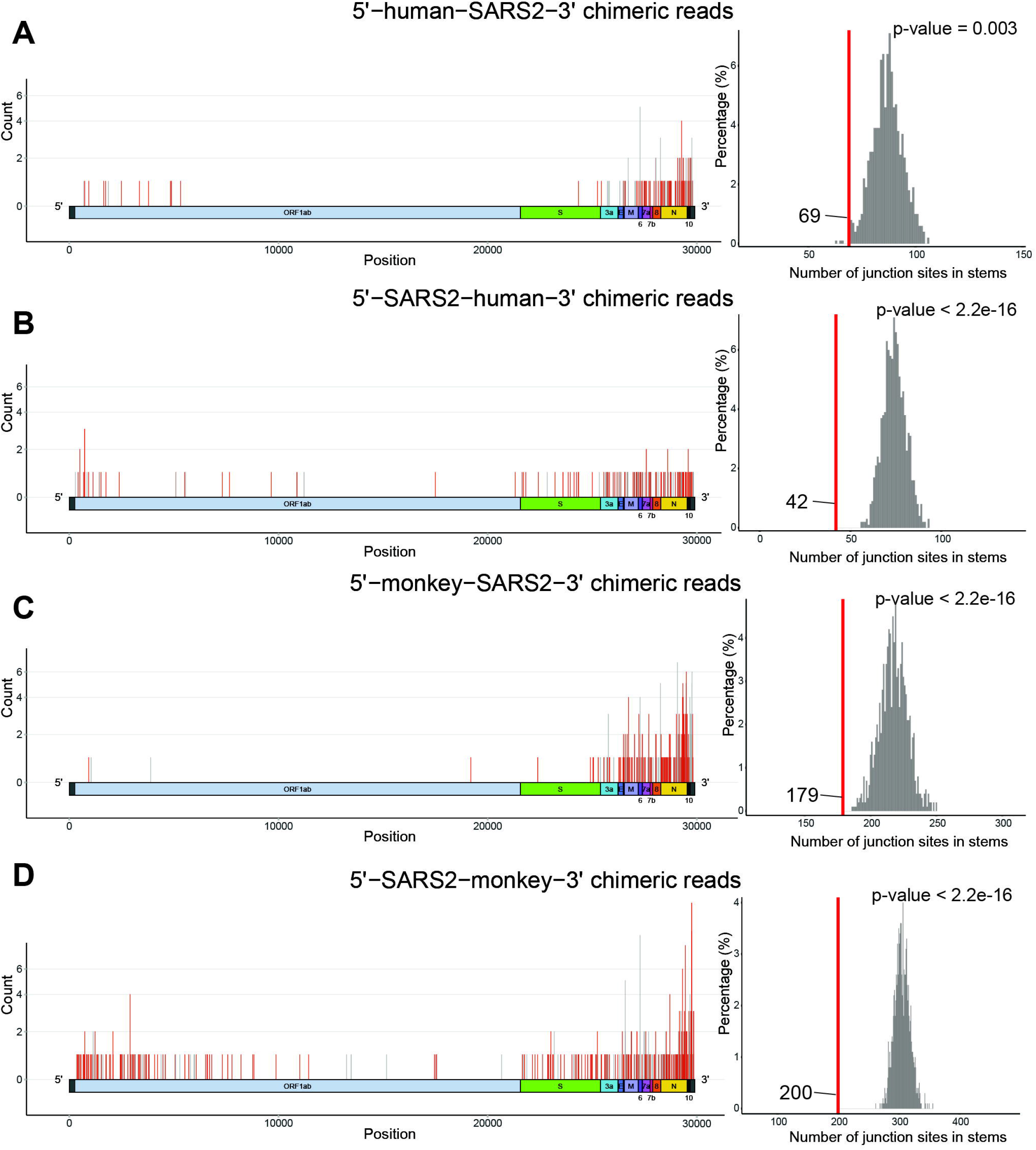
Locations of the chimeric read junction sites and permutation tests for the number of junction sites in stems. Diagrams show how frequently junction sites occur at each position on the SARS-CoV-2 genome for **(A)** 5’-human-SARS2-3’, **(B)** 5’-SARS2-human-3’, **(C)** 5’-monkey-SARS2-3’, and **(D)** 5’-SARS2-monkey-3’ chimeric reads. Positions are colored based on the secondary structure of the SARS-CoV-2 RNA, with red lines indicating that the position is in the non-stem region, while gray indicates that the position is located in the stem region. Histograms following each diagram show the corresponding results of permutation tests used to test if the junction sites of chimeric reads are within base-paired regions of the viral RNA. Each test consists of 1000 permutations and the actual frequency of junction sites occurring in the stem regions is marked with a vertical red line.

Previous studies have also found that indel formation and template switching events preferentially occur in the loops and stems formed in the RNA secondary structure (2, 3). First a permutation test was used to investigate if junction sites were commonly located in stems (positions that form base-pairs) or non-stem regions (non-base-paired positions) in the viral RNA. The results of this test showed a significant (P-values < 0.01) preference for the formation of junctions in non-base-paired regions of the RNA secondary structure (Fig.1). One-sided Fisher’s exact tests were performed to explore if junction sites were enriched in specific types of RNA structures. Consistent with the results of the permutation test, stems were under-represented at the junction sites (Supplementary Table 3). We speculate that the non-base-paired regions of the SARS-CoV-2 RNA may be more susceptible to stochastic template-switching events due to their more “open” configurations, where the viral RdRp could easily attach or detach as it moves along the RNA.

An examination of the types of human gene sequences found in the chimeric sequences revealed an enrichment of non-coding RNAs and highly expressed genes. We found that a disproportionate number of non-coding RNAs, mainly long non-coding RNAs (lncRNAs), were forming parts of the chimeric reads compared to their abundance in the human genomes. These non-coding RNA chimeric sequences made up 8.8% and 10.5% of the chimeric reads detected in the Caco and Calu cell lines, respectively, while non-coding sequences made up only 4% of the genes annotated in the human genome. This enrichment of non-coding RNA chimeric sequences was tested using Fisher’s exact test confirming that the trend was significant (Caco cells: odds ratio=2.2, P-value=0.043; Calu cells: odds ratio=2.8, P-value=0.001). When analyzed in the context of the expression level of the host genes in each sample, we also observed an enrichment for highly expressed genes forming parts of the chimeric sequences (Fig.2). This enrichment was confirmed through the Mann-Whitney U tests showing that the trend was significant in the two human cell lines (P-value < 2.2e-16 for both) and the *Chlorocebus sabaeus* (green monkey) cell line (P-value < 2.2e-16). These results appear to highlight two groups of sequences that are forming chimeric RNAs, structural RNAs like lncRNAs, which may be susceptible due to their secondary structures, and highly expressed genes, which would have more RNA molecules present for template-switching events to occur with. This suggests that the formation of chimeras is largely stochastic, with factors like the abundance of RNAs playing a large role, but that certain RNA molecules may be more susceptible to these events due to their structure.

**Fig.2.**
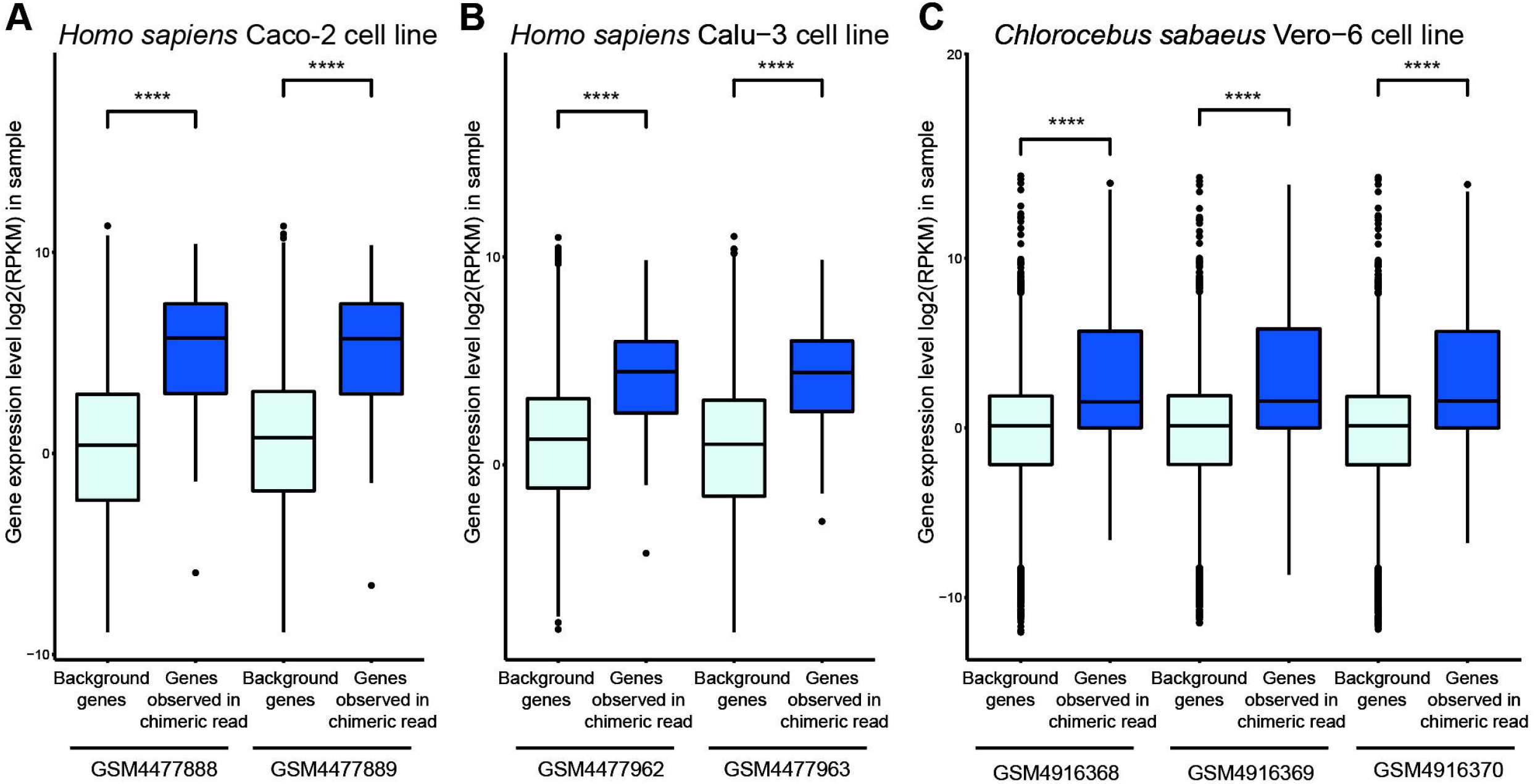
The expression level of host genes observed in chimeric reads. The expression level of host protein-coding genes observed in chimeric reads is significantly higher than the background protein-coding gene expression level based on studies on **(A)** *Homo sapiens* Caco-2 cell line, **(B)** *Homo sapiens* Calu-3 cell line, and **(C)** *Chlorocebus sabaeus* Vero-6 cell line.

### Systematic search for host-derived insertions in SARS-CoV-2 genomes

We performed a survey of the GISAID SARS-CoV-2 genomes to identify insertions with potential host origins. Insertions were detected based on alignments and comparison to the Wuhan-Hu-1/2019 reference genome. Only insertions greater than or equal to 21 nucleotides long and that were found outside of the 5’ and 3’ untranslated regions were considered in subsequent analyses (Supplementary Table 4). Of the 36 insertions that were found, 17 of them were found in multiple SARS-CoV-2 genomes but were not monophyletic. Upon further examination, the genomes containing these insertions tended to be sequenced by the same labs around the same times making it likely that these detected insertions are due to library preparation or sequencing errors rather than the result of multiple independent insertion events in different viral lineages. Of the 19 other insertions, 16 of them were only detected in a single genome, and while many of these had plausible hits to human genes, it is difficult to assess if these are true insertions or library preparation or sequencing artifacts due to their limited presence.

The three remaining insertions were from monophyletic virus variants and were further examined to determine if they had plausible homologous sequences in the human genome. Two of the insertions were found to be identical to conserved segments of the 28S and 18S rRNAs and were analyzed further. The remaining insertion was 21 nucleotides long and was found in 6 SARS-CoV-2 genomes of the Alpha B.1.1.7 lineage. These genomes were collected in early March of 2021 from England, United Kingdom by two laboratories, and sequenced at the same location using the same sequencing platform. The raw reads were available for two of the genomes, namely England/ALDP-13C8C28/2021 (EPI_ISL_1331302) and England/QEUH-13C1955/2021 (EPI_ISL_1332461), and were examined directly, providing confirmation that the insertion was present and likely not an artifact. Unfortunately, no plausible source for this insertion was able to be identified using a BLAST search in the NCBI non-redundant nucleotide database and a collection of coronavirus genomes with a cutoff E-value of 1e-2, and it was not analyzed further.

### 28S rRNA-derived insertion in SARS-CoV-2 genomes

We detected a 27-nucleotide long insertion in five SARS-CoV-2 genomes (Supplementary Table 5 and Fig.3A) at position 7120 of the reference genome (China/Wuhan-Hu-1/2019). The five genomes containing the 28S rRNA-derived insertions were collected by different laboratories and were sequenced on different sequencing platforms, making it extremely unlikely that laboratory error is responsible for the presence of the insertions. The five genomes belong to a monophyletic group. In this clade, there are three other variants whose assembled genomes do not contain the insertion. We were able to obtain access to the raw genome sequencing data of two of the three variants — USA/WA-PHL-005726/2021 (EPI_ISL_6259191) and USA/HI-H215617/2021 (EPI_ISL_6540096). We then did further analysis on the raw sequencing to check if the insertion is indeed missing. First, we generated consensus genome sequences based on the alignment of sequencing reads to the SARS-CoV-2 reference genome and found the consensus sequences did not contain the insertion. Next, we manually added the 28S rRNA-derived insertion at position 7120 of the consensus genome and compared the reads exclusively aligned to the consensus genome with the insertion and the reads exclusively aligned to the consensus genome without the insertion. We found that 99.76% (8700/8721 for EPI_ISL_6259191) and 99.93% (1502/1503 for EPI_ISL_6540096) of the exclusively-mapped reads support the presence of the insertion in the genomes. The reason that the insertion is missing in the submitted genomes (EPI_ISL_6259191 and EPI_ISL_6540096) is likely that the assembly was generated using an insertion-unaware approach, such as reference-based consensus calling. For the only one variant that was missing the insertion in the genomes, we are not able to assess if it is due to failure to identify the insertion based on the consensus caller or the subsequent loss of the inserted sequence.

**Fig.3.**
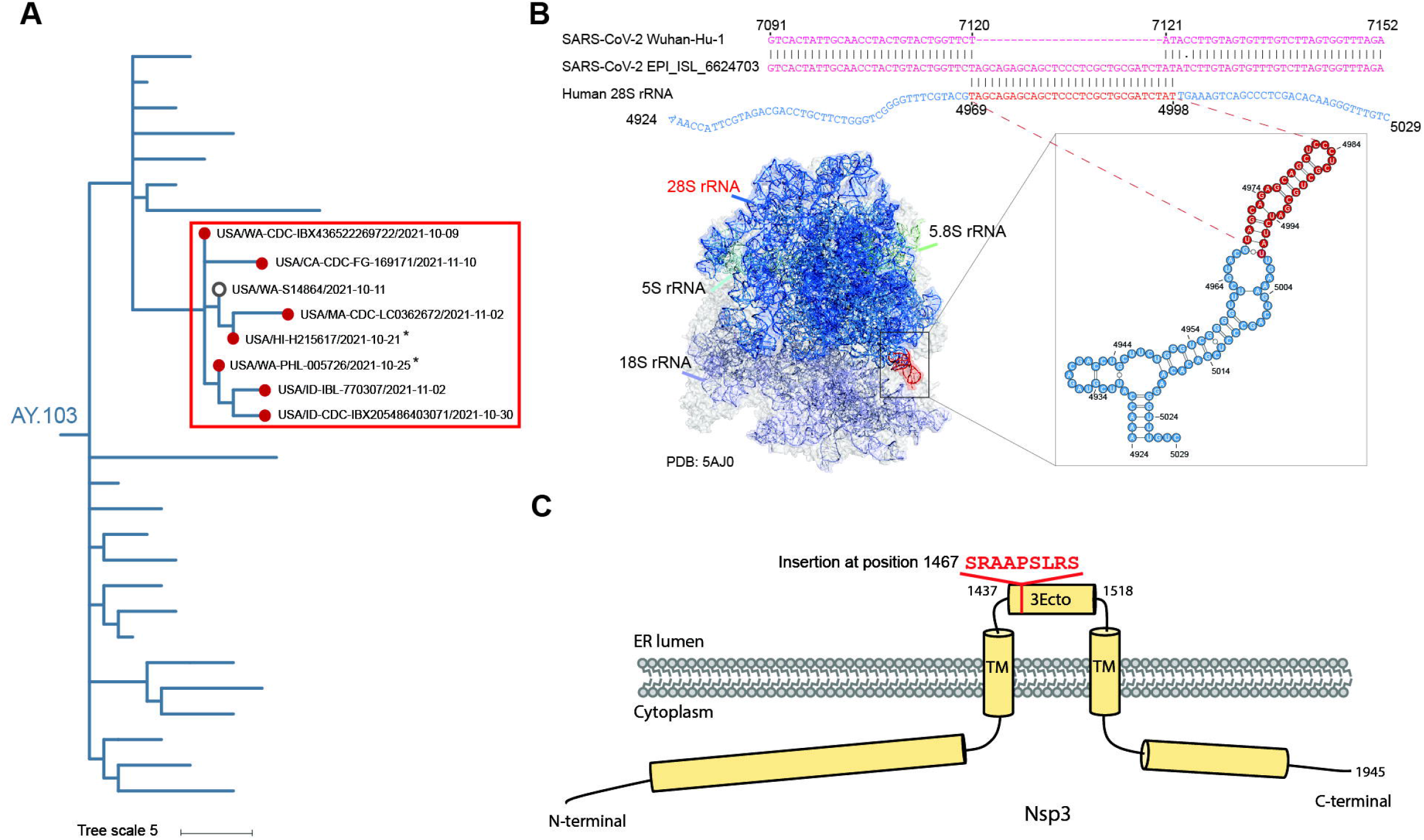
The 28S rRNA-derived insertion in SARS-CoV-2 genomes. **(A)** The phylogeny tree shows the genomes containing the human 28S-derived insertion. The clade where the insertion was detected is highlighted with a red box and the genomes with the insertion are marked with red circles at the tips. The asterisk (*) indicates that the insertion should be present in the variant based on raw sequencing data. **(B)** The insertion in SARS-CoV-2 genomes potentially originate from the host 28S rRNA shown by the sequence alignment of SARS-CoV-2 reference genome (NCBI accession: NC_045512.2, GISAID accession: China/Wuhan-Hu-1/2019) (pink), USA/CA-CDC-FG-169171/2021 (NCBI accession: OL591909.1, GISAID accession: EPI_ISL_6624703) (pink) and human 28S rRNA (chain A2 of PDB 5AJ0) (blue). There are five possible alignments for mapping this insertion to the reference. Only the alignment with the sequence inserted after the 3^rd^ position of 2285^th^ codon in ORF1ab is shown. The putative insertion origin is colored in red. The numbers listed above and below the alignment indicate the positions of aligned bases in the original sequences. The insertion sequence (red) was mapped to the 28s rRNA (blue) in a human polysome 3D structure (PDB: 5AJ0). A zoom-in view of the RNA secondary structure shows that the insertion is located on the No. 94 stem of domain 7 (position: 4969-4998) 28S rRNA region (highlighted red). **(C)** Diagram shows the position of the human 28S rRNA-derived insertion in the ectodomain (3Ecto) of Nsp3 protein.

By performing the BLAST search for this insertion against the human transcripts (Release 109 RNAs), an exact match (E-value: 2e-06) of this insertion was found in the nucleotide sequences of 28S ribosomal RNA (Fig.3B). We observed an extra three overlapping bases in the pairwise alignment of SARS-CoV-2 variants containing the insertion and the human 28S rRNA sequence, extending the length of identity nucleotide bases from 27 nucleotides to 30 nucleotides. The identical region was located at positions 4969-4998 of the human 28S rRNA (based on the structure of PDB 5AJ0 Chain A2) and makes up part of the highly conserved loop 94 stem of domain 7 of the rRNA molecule according to the Gorski et al.’s segmentation of human 28S rRNA (26) (Fig.3B).

Due to the high level of sequence conservation of 28S rRNA, asserting the origin of the insertion-related 30 nucleotide sequences is impossible based on sequence identity alone. In the human genome (GRCh38 release 105), three 28S rRNA gene copies in chromosome 21 and one copy in chromosome 12 contain the exact 30 nucleotide sequences. When we searched the 30 nucleotide sequences in the LSU rRNA database downloaded from SILVA (27), 98 organisms were found to contain the sequences. The last common ancestor of these 98 organisms is *Euteleostomi* (bony vertebrates). Given the fact that the insertion emerged from the SARS-CoV-2 variant circulating in humans, the originating organism of the 28S rRNA-derived insertion is most likely humans.

The nine amino acid insertion is located at position 1467 of the ectodomain (3Ecto) in the Nsp3 protein, the only domain of this protein located on the lumenal side of the endoplasmic reticulum (Fig.3C). Nsp3 along with Nsp4 and Nsp6 have been shown to be involved in the formation of double membrane vesicles in coronavirus infected cells (28, 29). The 3Ecto domain is specifically involved in the recruitment of Nsp4 and has been shown to be an essential component of Nsp3 for correct double-membrane vesicle formation (28). At this point, it is unclear if this insertion would have had an effect on viral fitness, but given its location in the 3Ecto domain, it is possible that the insertion could have an effect on the interactions between Nsp3 and other proteins and on the membrane rearrangement process.

The monophyletic group with the 28S rRNA-derived insertion belonged to the AY.103 group of the delta lineage (30) (Fig.3A). The AY.103 variant was first detected worldwide on January 1^st^, 2021 and in the USA on January 2^nd^, 2021. The clade containing the 28S rRNA-derived insertion is defined by five nucleotide mutations (T7900C, A10420T, C18646T, C25721T, and C29668T). By September 2021, AY.103 had become the most common delta lineage in the United States and has continued to be responsible for a significant fraction of cases until the recent emergence of the Omicron variant (31). The five genomes containing the 28S rRNA-derived insertion were collected between October 9^th^ and November 10^th^ in 2021 from the states of Washington, Idaho, Massachusetts, and California, indicating that these variants were likely being transmitted over this timeframe, but the extent to which it was being spread seems to be low as Idaho was the only state where multiple genomes were collected from and no genomes containing the insertion have been reported since. Based on the limited spread of the viruses containing the 28S rRNA-derived insertion, it is likely that the insertion might not confer phenotypic advantages or is possibly disadvantageous to the virus. Nonetheless, our data show that AY.103 lineages containing this insertion were viable and were transmitted for a short period of time.

### 18S rRNA-derived insertion in SARS-CoV-2 genomes

A 24-nucleotide insertion was detected in two genomes at position 27492 in the genome of the reference genome (China/Wuhan-Hu-1/2019) (Supplementary Table 5). A sequence search against human transcripts (Release 109) was performed using BLAST (32), resulting in the identification of an exact match to a 24 nucleotide stretch (E-value: 2e-5) of the 18S rRNA sequence. When aligned to the full 18S rRNA sequence, it was found that the identical region extended one additional nucleotide outside of the insertion region, bringing the identical stretch to 25 nucleotides (Fig.4A). The insertion was identical to a highly conserved region of the 18S rRNA (at positions 399-423 in 18S rRNA), consisting of a portion of the helix 12 of the 5’ domain (33, 34). In the human genome alone there are five copies of the 18S rRNA gene on chromosome 21 that contain identical matches for this 25 nucleotide sequence. When compared to the SSU rRNA SILVA database (27), identical sequences were found in the 18S sequences of 2289 organisms, which had a common ancestor of *Opisthokonta* (Fungi/Metazoa group). Considering that the viral samples were circulating in human populations, it is highly likely that the insertion was derived from human 18S rRNA.

**Fig.4.**
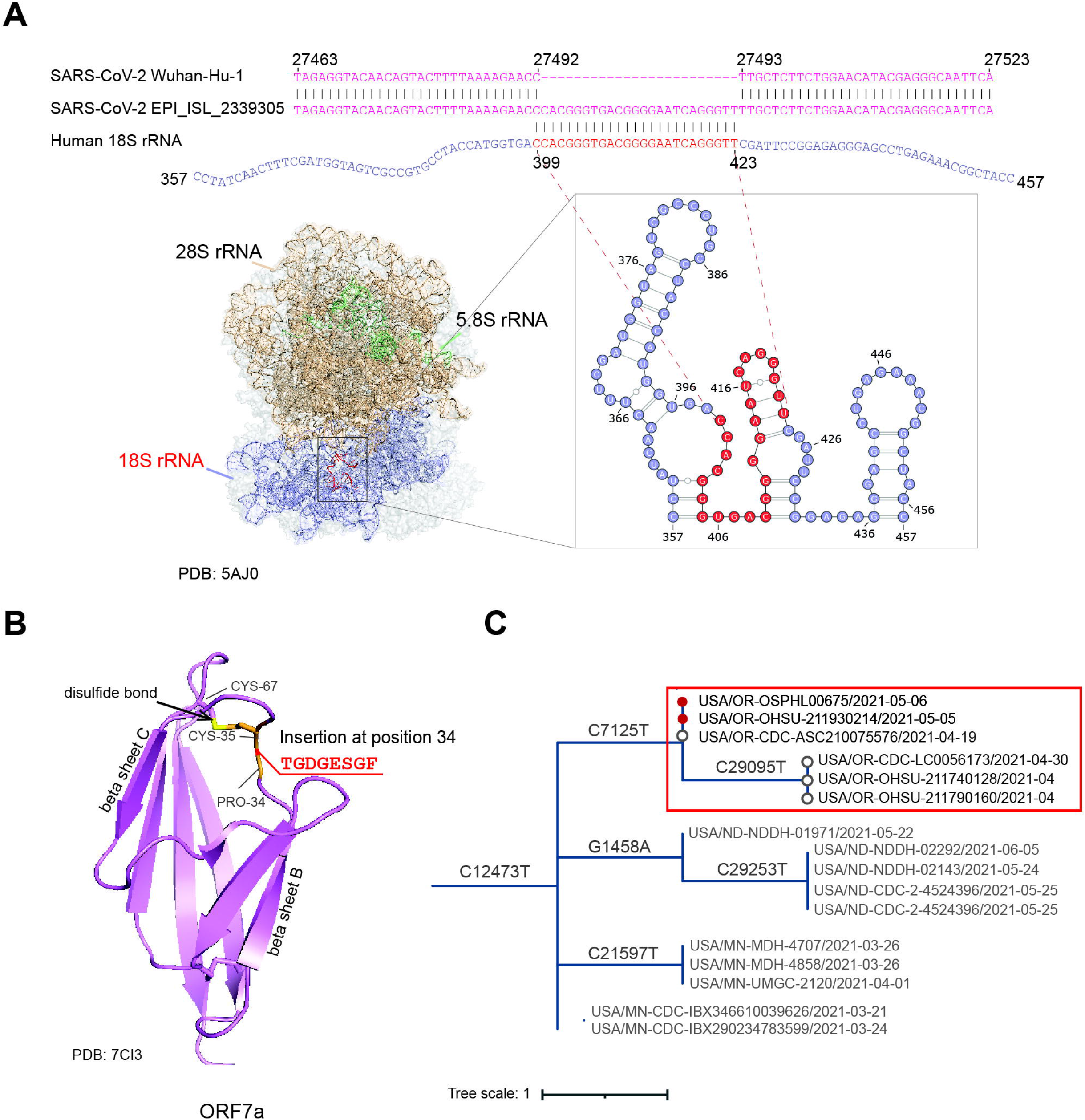
The 18S rRNA-derived insertion in SARS-CoV-2 genomes. **(A)** The insertion in SARS-CoV-2 genomes potentially originates from the host 18S rRNA shown by the sequence alignment of SARS-CoV-2 reference genome (NCBI accession: NC_045512.2, GISAID accession: China/Wuhan-Hu-1/2019) (pink), USA/OR-OSPHL00675/2021 (GISAID accession: EPI_ISL_2339305) (pink) and human 18S rRNA (purple). The putative insertion origin is colored in red. The numbers listed above and below the alignment indicate the positions of aligned bases in the original sequences. The insertion sequence (red) was mapped to the 18S rRNA (purple) in a human polysome 3D structure (PDB: 5AJ0). A zoom-in view of the RNA secondary structure shows that the insertion covers parts of helices 11 and 12 of the 5’ domain of the 18S rRNA. The location of the putative insertion sequence is highlighted red. **(B)** Diagram shows the position of the human 18S-derived insertion on the structure of the SARS-CoV-2 ORF7a protein (PDB: 7CI3). **(C)** The phylogeny tree shows the genomes containing the human 18S-derived insertion. The clade where the insertion was detected is highlighted with a red box and the genomes with the insertion are marked with red circles.

The insertion is in the SARS-CoV-2 ORF7a protein, encoding an eight amino acid sequence that is located between the proline and cysteine at positions 34 and 35 in the reference protein sequence (Fig.4B). The cysteine at position 35 is known to form a disulfide bond with a cysteine at position 67 and is thought to help stabilize the beta-sheet structure (35, 36) and the possible functions of the proline at position 34 are not known. The ORF7a protein has been shown to contain an immunoglobulin-like ectodomain between residues 16 and 96 on the protein which is thought to have a role of binding to human immune cells and modulating immune response (35–37). Given the proximity of the insert to the disulfide bond forming cysteine at position 34 and the size of the insert it is possible that this insert would have an effect on the overall structure and immunoregulatory functions of ORF7a, but without additional evidence, the effect of this insertion on the fitness of the virus remains unknown.

The two genomes containing the 18S rRNA insertion were from the same clade in the Alpha B.1.1.7 SARS-CoV-2 lineage, which was first identified in England, United Kingdom in mid-December of 2020 (Fig. 4C). This variant was designated as a variant of concern due to its transmissibility and large number of mutations and quickly became the dominant variant in England while spreading to other countries (38). The genomes containing the 18S rRNA-derived insertion, along with the other four genes in the same clade, were collected in April and May of 2021 in Oregon, United States. The genomes from the variants containing the insertion were collected and sequenced by different labs using different sequencing platforms, making it unlikely that the insertion was a sequencing or library preparation artifact. We did not detect the insertion in any of the other four genomes from this clade, indicating that either they do not have the insertion, they have it but it was not detected, or that the insertion was only acquired in a sub-clade within this group. After May of 2021, no new genomes containing this insertion were collected, indicating that the period during which these lineages were circulating may have been brief. While these viral variants seem to be viable and transmitted for a short period of time, the insertion likely does not confer a significant advantage or may be disadvantageous for the virus resulting in its limited spread.

## DISCUSSION

Insertions in the SARS-CoV-2 genome can be introduced through multiple mechanisms and have the potential to give rise to new variants with enhanced infectivity, pathogenicity, and antibody escape (2, 6), but the source of these insertions is often difficult to determine and has been hotly debated (5, 7). Leveraging available direct RNA sequencing data and an analysis of SARS-CoV-2 genomes, we have found evidence of the formation of viral-host chimeric RNA sequences and described two novel human-derived genomic insertions present in circulating variants of SARS-CoV-2.

Through our screening of direct RNA-seq data from SARS-CoV-2 infected cell lines, we found that viral-host chimeric RNAs were rare but were present in approximately half of the samples analyzed. The chimeric reads all contained positive-sense viral RNA sequences, indicating that these chimeric sequences are not the result of the integration of the viral genetic material into the host genome, which would have resulted in a nearly equal mix of positive and negative sense viral sequences (17). This process does appear to be stochastic in nature though, with no preference for starting with host or viral sequences during chimera formation and a higher frequency of chimeras being formed with highly expressed genes in the cells. The regions in the RNA where these template switching events occur appears to be influenced by the secondary structure of the viral RNA, possibly due to certain structures being more susceptible to template switching events similar to what has been reported in previous studies (2, 3). The accurate determination of the exact junction boundaries and potential base-pairings were hindered by the high error rate of 14% in direct RNA-sequencing data and the limited number of host-viral chimeras detected in this study. The exact molecular basis for the viral-host chimera remains unclear and future investigation with larger sets of error-corrected direct RNA-seq data of SARS-CoV-2 could be beneficial to address this question.

The formation of host-viral chimeric mRNAs or subgenomic RNAs could mostly be transient events, not having a long-term impact on viral fitness, but the possibility of human-derived insertions in the coronavirus genomes could have significant implications considering the role that genomic insertions seem to have in the evolution of new SARS-CoV-2 variants (5, 6). The putative 18S and 28S-derived insertions were identified in circulating variants of the SARS-CoV-2, and while these particular variants did not seem to spread widely, they do provide evidence that human genetic material can be a source of genomic insertions in SARS-CoV-2. Interestingly, rRNAs have been established to be a source of insertions in influenza genomes, in some cases resulting in significantly more pathogenic viral variants (39, 40). It has been speculated that these recombination events often occur with host rRNAs due to their abundance in the cells, the presence of recombination hotspots on rRNA molecules, and the utilization of host rRNAs during viral replication (39). Similar factors may play a role in the formation of these rRNA-derived insertions in SARS-CoV-2, but the formation of double-membrane vesicles during SARS-CoV-2 would seemingly complicate this process. There may be accidental capture of host RNAs inside of the double-membrane vesicles during their formation or some crossover of host RNA from the cytosol, but evidence of this is lacking and warrants further investigation.

## CONCLUSIONS

Overall, our results suggest that viral-host chimeric sequences can be formed, likely through stochastic RdRp template switching events. Furthermore, we have identified two long insertions in SARS-CoV-2 genomes in previously circulating variants which are likely derived from human ribosomal RNAs. While the source of smaller insertions that are present in many SARS-CoV-2 genomes are still difficult to identify due to their short lengths, these results provide evidence that bolsters the hypothesis that some of them are derived from human genetic material. The mechanisms at work in the formation of these chimeric RNAs and genomic insertions are still unclear but warrant further study considering the potential importance of these processes in viral evolution and the emergence of new variants.

## METHODS

### Identification of host-virus chimeric reads in SARS-CoV-2 direct-RNA seq data

The nanopore direct RNA-seq data from SARS-CoV-2 infected cell lines were downloaded from the NCBI SRA database (Supplementary Table 1). All reads were quality trimmed using NanoFilt v2.8.0 (41), to remove the first 50 nucleotides of each read and require an average quality score of at least 10 over the length of the read. The trimmed reads were then mapped using Minimap2 v2.23 (42) to the SARS-CoV-2 reference genome (NCBI GenBank accession: NC_045512.2) (43), and either a reference *Chlorocebus sabaeus* transcriptome (ftp://ftp.ensembl.org/pub/release-105/fasta/chlorocebus_sabaeus/) or human transcriptome (ftp://ftp.ensembl.org/pub/release-105/fasta/homo_sapiens/). The mapping files were converted to the Pairwise mApping Format (PAF) using the paftools script that is part of Minimap2 (42). Reads that mapped to both the host and SARS-CoV-2 transcriptomes were extracted for analysis as potential chimeric sequences. To avoid including chimeric reads that resulted from technical artifacts such as those caused by misinterpretation of open-pore states by base-calling softwares (19), additional quality filtering was applied to the chimeric reads. The distance between the mapped regions of the virus and the host sequence on the chimeric reads was required to be less than 15 nucleotides, the junction was required to be formed in the middle of the genes (not within the last 50 nucleotides of the first gene sequence, nor the first 50 nucleotides of the second gene sequence), and the quality score within 20 bp of either side of the junction was required to be higher than the 20th percentile quality score for that read.

### Mapping short reads to direct-RNA seq chimeric reads

We collected paired-end sequencing data on five samples with corresponding direct RNA-sequencing data. The short reads were first preprocessed with fastp v0.23.1 (44) and then mapped to the chimeric reads from the same samples by using Minimap2 v2.23 (42) with options “-ax sr -w 5” to tolerate the high error rate of the Nanopore direct RNA-sequencing reads (45). Read pairs spanning the junctions were detected and counted with a custom script. The numbers of read pairs supporting the chimeric reads are provided in Supplementary Table 2.

### Analysis of junction positions in relation to viral RNA secondary structure

The RNA secondary structure of the SARS-CoV-2 reference genome was obtained from previous studies (46, 47) and bpRNA (48) was used to assign each residue to secondary structure elements. A junction site was considered in the stem if the two flanking nucleotides were in the same stem. To investigate if junctions tend to happen in non-stem regions, the number of junctions occurring in base-paired positions were calculated and compared with a background distribution for the numbers of junctions located in stems derived from a 1000-time random sampling of the same number of sites along the viral RNA strand. To further examine which types of structural elements are over- or under-presented at junction sites in virus-host chimeric reads and in host-virus chimeric reads, one-sided Fisher’s exact test was performed.

### Analysis of the expression level of host genes observed in chimeric reads

Gene expression profiles for two SARS-CoV-2 infected Caco-2 cell line samples (GSM4477888, GSM4477889), two SARS-CoV-2 infected Calu-3 cell line samples (GSM4477962, GSM4477963), and three SARS-CoV-2 infected Vero-6 cell line samples (GSM4916368, GSM4916369, GSM4916370) were downloaded from the GEO database. The read counts of each gene were normalized by the total number of reads in each sample and by the gene length (RPKM) to represent the gene expression level. The background gene set was composed of all expressed protein-coding genes in the cell line. To evaluate whether the expression level of the host protein-coding genes in chimeric reads is significantly greater than the expression level of the background gene set, a one-sided Mann-Whitney U test was performed for each sample.

### Identification of insertions in SARS-CoV-2 genomes

The SARS-CoV-2 genomes available at GISAID (https://www.gisaid.org/) on 2021-12-17 were downloaded for analysis (n=6,163,073). The sequences were then processed by NextClade CLI v1.7.0 (49) which generated a multiple sequence alignment against the reference genome (Wuhan-Hu-1/2019) and provided a list of single nucleotide polymorphisms, insertions, and deletions associated with each genome sequence. Only sequences that passed all quality controls and were assessed as “good” applied by NextClade were used for further analysis (n=5,226,229). Insertions greater than or equal to 21 nucleotides long and found outside of the 5’ and 3’ untranslated regions of the viral genomes were kept. They were searched in the NCBI non-redundant nucleotide database and a collection of coronavirus genomes with BLASTN (E-value≤1e-2) (32) to explore their possible origins.

### Monophyletic test

To check if the insertions of interest formed a monophyletic group, all genomes that contained the same insertion were analyzed using UShER: Ultrafast Sample placement on Existing tRee v0.5.1 (50) against a phylogenetic tree with available genomes (n=6,257,569) from GISAID, GenBank, COG-UK and CNCB generated by sarscov2phylo pipeline v.13-11-20 (51). The sequences are placed within an updated global subsampled SARS-CoV-2 phylogenetic tree and local subtrees are computed to show more sequences with the same context of the ones being analyzed.

### Verification of insertions with raw sequencing data

The raw genome sequencing data of USA/WA-PHL-005726/2021 (EPI_ISL_6259191) and USA/HI-H215617/2021 (EPI_ISL_6540096) were analyzed to check if the insertion is indeed missing. The raw sequencing reads were processed for quality control using fastp v0.23.1 (44) with default parameters and mapped to the SARS-CoV-2 reference genome using BWA mem v0.7.17 (52). Primer sequences in reads of EPI_ISL_6259191 were soft clipped using ivar trim (parameters: -m 1 -q 0 -s 4 -e) and reads in amplicons with variants in primer binding sites were removed by ivar removereads v1.3.1 (53). The sequencing data of EPI_ISL_6540096 were preprocessed by the providing laboratory and the primers were removed. Consensus genome sequences were generated based on the alignments and it was found that the consensus sequences did not contain the insertion. The 28S rRNA-derived insertion was manually added at position 7120 of the consensus genomes to generate consensus genomes with the insertion. The alignment files were converted to FASTQ format using samtools fastq command v1.14 (54) and re-aligned to the consensus genomes with or without the insertion using bowtie v2.4.4 (45) (parameter: --xeq). Reads exclusively aligned to the consensus genome with the insertion and exclusively aligned to the consensus genome without the insertion were identified with a custom script (https://github.com/ncbi/SARS2_host_derived_insertions/blob/main/verify_insertion/insertion_match_reads.py).

## Data availability

The datasets generated in this study and scripts are available in the github repository, https://github.com/ncbi/SARS2_host_derived_insertions.

## Authors’ contributions

YY was involved in the execution of the analyses, interpretation of the results, and writing and revision of the manuscript. KD was involved in the interpretation of the results and writing and revision of the manuscript. RF was involved in the execution of the analyses and writing of the manuscript. XJ was involved in the conceptualization, planning, interpretation of the results, and revision of the manuscript. All authors read and approved the final manuscript.

## Acknowledgments

This work utilized the computational resources of the NIH HPC Biowulf cluster (http://hpc.nih.gov). We would like to thank Eugene V. Koonin and Sofya K. Garushyants for their thoughtful comments on our manuscript and their code on how to perform permutation tests and plot the results provided at https://github.com/garushyants/covid_insertions_paper. We gratefully acknowledge the researchers from the originating laboratories responsible for obtaining the specimens and the submitting laboratories where genetic sequence data were generated and shared via the GISAID Initiative, on which this research is based (Supplementary Table 6). We want to particularly thank the Washington State Public Health Laboratories and the State of Hawaii Laboratories Division for sharing the raw sequencing data for the genomes USA/WA-PHL-005726/2021 and USA/HI-H215617/2021.

## SUPPLEMENTAL MATERIAL

### Supplementary Tables

Supplementary Table 1: Direct RNA-seq data analysis. Metadata associated with all 30 of the analyzed directed RNA-seq samples is provided along with the number of reads mapped to the SARS-CoV-2 transcriptome and the count and frequency of chimeric reads in each sample. The counts of chimeric reads in the host to virus and virus to host orientation are listed for each sample. The references DOI for each sample is also listed.

Supplementary Table 2: Chimeric reads supported by spanning junction short reads.

Supplementary Table 3: Secondary structure element enrichment analysis of nucleotide sequences at junction sites.

Supplementary Table 4: Long insertions identified in GISAID that are not derived from SARS-CoV-2. The location on the SARS-CoV-2 reference genome, insertion sequence, insertion length, and what gene they are located in are provided for each of the 36 detected insertions. SARS-CoV-2 genomes with the insertion, whether those genomes were monophyletic, and short descriptions of putative matches are also provided for each insertion.

Supplementary Table 5: Information on genomes related to the three verified insertions. Metadata associated with each of the SARS-CoV-2 genomes with putative insertions that were analyzed including the variant types, collection dates, collecting labs, and sequencing methods are provided.

Supplementary Table 6: GISAID acknowledgement table.

